# Searching deeply into the conformational space of glycoprotein hormone receptors. Molecular dynamics of the human follitropin and lutropin receptors within a bilayer of (SDPC) poly-unsaturated lipids

**DOI:** 10.1101/2023.08.09.552573

**Authors:** Eduardo Jardón-Valadez, Alfredo Ulloa-Aguirre

## Abstract

Glycoprotein receptors are a subfamily of G-protein coupled receptors, including the follicle hormone (FSH) receptor (FSHR), thyroid-stimulating hormone receptor (TSH), and luteinizing/chorionic gonadotrophin hormone receptor (LHCGR). These receptors display common structural features such as a prominent extracellular domain, with a leucine-rich repeats (LRR) stabilized by β-sheets, a long and flexible loop known as the hinge region (HR), and the transmembrane (TM) domain with seven α−helices interconnected by intra- and extracellular loops. Binding of the ligand to the LRR resembles a hand coupling transversally to the α− and β−subunits of the hormone, with the thumb being the HR. The structure of the complex of FSHR-FSH suggests an activation mechanism in which Y335 at the HR binds into a pocket between the α− and β−chains of the hormone, leading to an adjustment of the extracellular loops. In this study, we performed molecular dynamics (MD) simulations to identify the conformational changes for the FSHR and LHCGR. We set up an FSHR structure as predicted by AlphaFold (AF-P23945); for the LHCGR structure we took the cryo-electron microscopy structure for the active state (PDB:7FII) as initial coordinates. Specifically, the flexibility of the HR domain and the correlated motions of the RLL and TMD were analyzed. From the conformational changes of the LRR, TMD, and HR we explored the conformational landscape by means of MD trajectories in all-atom approximation, including a membrane of polyunsaturated phospholipids. The distances and procedures here defined may be useful to propose reaction coordinates to describe diverse processes such as the active-to-inactive transition, to identify intermediaries suited for allosteric regulation, and biased binding to cellular transducers in a selective activation strategy.

**Author summary:** In the present study, we describe the results from a computational microscopy perspective (also known as molecular dynamics simulation) at the atomistic resolution for the two gonadotropin hormone receptors, the follicle-stimulant hormone receptor and the luteinizing/chorionic gonadotropin hormone receptor, which are essential for reproduction in humans. Several dysfunctional mutations in these receptors, leading to reproductive failure, have been detected in the clinical arena. To better understand the process whereby these two receptors perform their signaling tasks, triggering an intracellular response upon binding of their cognate agonist at the extracellular side, we assembled the receptor structures in a membrane bilayer of phospholipids with water molecules as solvent at both sides of the membrane. The systems included nearly 200 thousand atoms, each moving around at 300 kelvin and 1 bar given the interactions (attractive or repulsive forces) from each other. As the motion equations are solved in each time step (at femtoseconds time scale), the system evolves over time during hundreds of nanoseconds (millions of time steps) for three independent replicates. The receptor conformation, therefore, may display non-random motions due to the stability of specific structures in the complex molecular environment, including the hydrophobic membrane core, the bilayer interfaces, and the aqueous medium. From analysis of simulation trajectories and structural changes of the receptors, we could identify the main conformational changes exhibited by each receptor explored in a model cellular environment. We discussed the roll of the hinge domain at the extracellular domain in triggering the receptor conformational changes, as well as differences in the dynamics between these receptors in terms of the flexibility of the structures. Importantly, we proposed relative distances among the different receptor domains as parameters to characterize conformational intermediaries along a transition of states. Understanding of the signaling process in gonadotropin hormone receptors could be useful to explore new strategies for the modulation of the receptor functions, the bias of signaling pathways, or the selective binding of agonists.

## INTRODUCCIÓN

G protein-coupled receptors (GPCRs) are a large and functionally diverse superfamily of plasma membrane receptors that respond to a widely variable of endogenous and exogenous stimuli of diverse structures, from photons, odorants, and ions to lipids, neurotransmitters, peptide hormones and complex protein hormones. G protein-coupled receptors consist of single polypeptide chains of variable lengths which traverse the lipid bilayer seven times, forming characteristic transmembrane (TM) α-helices, connected by alternating extracellular and intracellular loops, with an extracellular NH_2_-terminus or ectodomain (ECD) and an intracellular COOH-terminus or Ctail [1, 2]. These membrane receptors currently represent an important therapeutic target for several diseases in humans; in fact, ∼30-40% of GPCRs of approved drugs target this family of membrane receptors [3, 4].

The receptors for the glycoprotein hormones (GPHR) [follicle-stimulating hormone (FSH) receptor or follitropin receptor (FSHR), lutropin receptor or luteinizing hormone/chorionic gonadotropin (LH and CG, respectively) receptor (LHCGR), and thyrotropin receptor or thyroid stimulating hormone (TSH) receptor (TSHR)] belong to a conserved subfamily of a GPCR family, the so-called Rhodopsin-like family, and more specifically, to the δ-group of this large class of GPCRs [1]. Structural features of the GPHRs include a large extracellular NH_2_-terminus, where recognition and binding of their corresponding ligands (FSH, LH and CG, or TSH) occur. This ECD exhibits a central structural motif of imperfect leucine-rich repeats (LRR; 12 in the FSHR; 8-9 in the LHCGR; 10 in the TSHR) [5–8], that is shared with other plasma membrane receptors involved in selectivity for ligand and specific protein-to-protein interactions (Fig. 1A)[9]. The carboxyl-terminal end of the ECD exhibits a domain critical for all GPHRs function: the so-called hinge region (HR); this particular region is involved in high affinity hormone binding, receptor activation and intramolecular signal transduction, and/or silencing of basal receptor activity in the absence of a ligand [10].

**Fig. 1.**
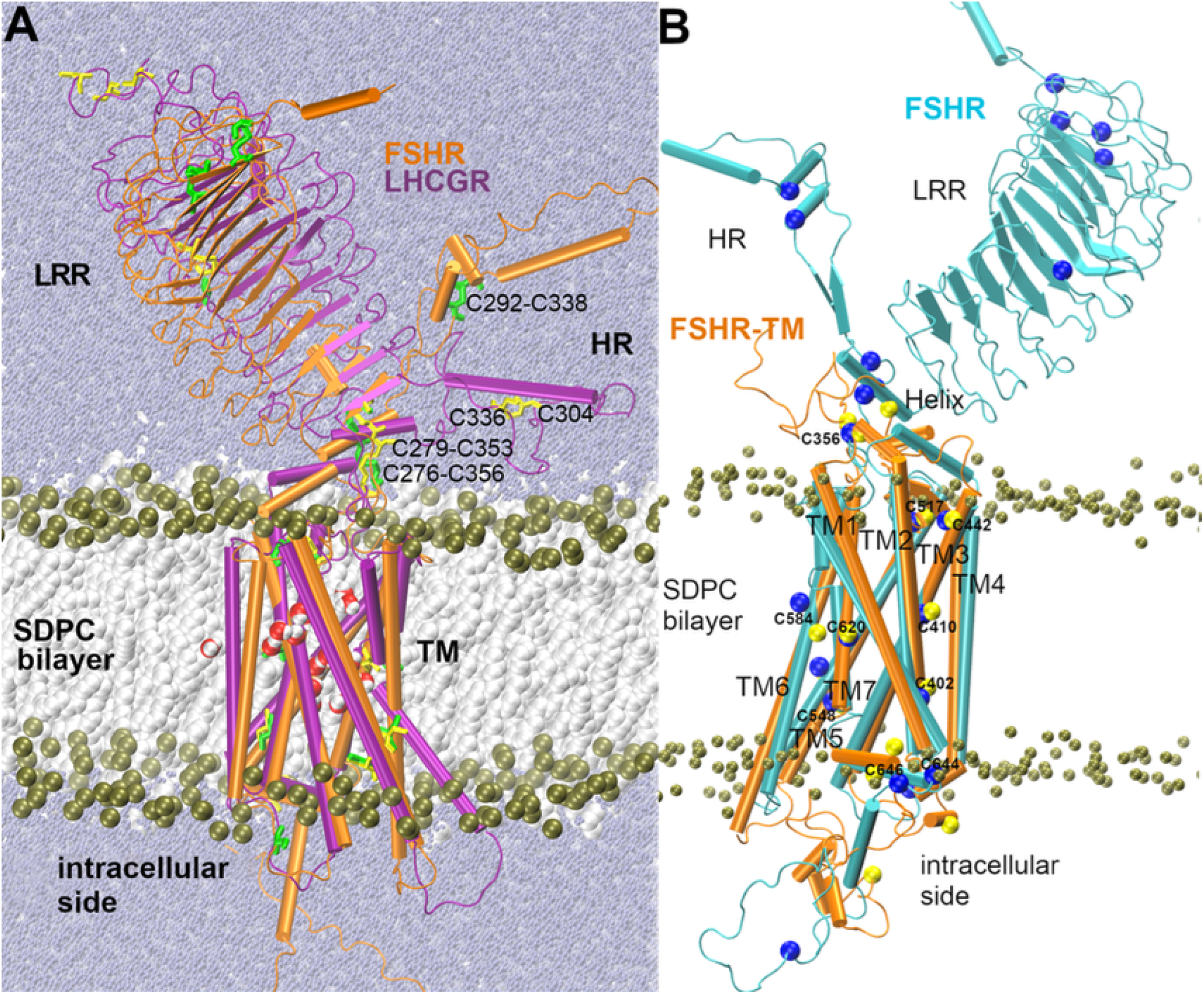
**A.** Simulation boxes for the FSHR (orange structure) and the LHCGR (purple structure). The leucine rich region (LRR), hinge region (HR) and transmembrane (TM) domain are indicated. The SDPC bilayer is depicted with the lipid tails as white spheres, and the phosphorus atoms of the lipid heads as olive green spheres. Solvent water molecules are depicted as small spheres in blue purple. Side chains of cysteine residues are depicted in licorice, green for the FSHR and yellow for the LHCGR. Disulfide bonds are identified for C292-C338, C276-C356 for FSHR, and C279-C336 of LHCGR. The bond C292-C338 at the HR of FSHR was not defined between C336 and C304, which is the homologous positions in the LHCGR. **B**. Structural alignment between FSHR (cyan structure) and the FSHR-TM (orange structure); for reference, Cα of cysteine residues are depicted as spheres in yellow and in blue for the FSHR-TM and FSHR, respectively. The open intracellular conformation suggests that the models corresponds to the activated conformation. Models were developed using different methods and the agreement provided an additional criterium for validation of the previously reported model [15].

The gonadotropin receptors (FSHR and LHCGR) play a critical function in reproductive function. They regulate spermatogenesis and ovarian follicular maturation (FSHR) as well as steroidogenesis (testicular Leydig cells, ovarian follicle, and placenta) as well as ovulation (LHCGR) [7, 11–13]. We have been interested in analyzing the effects of point mutations on function and tridimensional structure of the FSHR [7, 12, 14]. We have particularly focused on the understanding of the response of these receptors to their cognate agonists departing from changes in the conformational dynamics related to the biological function in dysfunctional variants [14, 15]. In principle, an extracellular signal is transmitted when the receptor is stabilized in an active conformation, which allows activation of the intracellular transducers coupled to the receptor. Nevertheless, the transition processes among different conformational states of the receptor triggered by the agonist are still incompletely understood [16, 17]. For example, the FSHR may activate several intracellular signaling cascades in function of coupling to distinct pathways mediated by several kinases and/or β−arrestins, that is to say, the receptors are more than simple binary interruptors, but rather function as allosteric microprocessors that respond to the ligand stimulus with different affinities to distinct transducers: a given ligand may favor activation of a particular signal transducer and, in turn, the transducer may increase the affinity of the receptor to the ligand [18].

Departing from the elucidation of the LHCG receptor in the active and inactive state by cryo-electron microscopy (3.5 Å), it is possible to identify the structural features of the GPHR subfamily [19]. The large extracellular ECD encompass the LRR, similar to a boxing glove, with the thumb formed by the HR. The α-helix (P272-N280) and the loop P10 (F350-Y359) of the HR conform a communication interface between the LRR and the TM domain. For example, the α-helix Q425-T435 of the EL1 and P10 exhibit interactions in the inactivating S277I and activating E254K mutations, suggesting that the positions K354 and K605 are important regulators of the activation or inhibition of the receptor [20]. On the other hand, the C304-C353 disulfide bond was not observed in the structure, in contrast to the C292-C338 bond in FSHR (Fig. 1A). Due to the absence of a crystal structure of the FSHR, our group has proposed a model for this receptor considering only the TM domain (FSHR-TM) [15, 21]. From the structural alignments of the LHCGR *vs* FSHR and FSHR *vs* FSHR-TMD, it is possible to identify that our FSHR- TM model corresponds to the active state of the FSHR (Fig. 1B). Among other findings derived from the trajectory analyses of molecular simulations of the FSHR and D408Y and I423T inactivating mutants, it was identified that helix 2 of the TM domain is an important communication hub for the propagation of intracellular signaling [14, 15].

As a continuation of previous studies [14, 15] in the present work we performed all-atom simulations for a FSHR model, but now including the corresponding LRR, HR and TM, as disclosed by the AlphaFold server (AF2; Fig. 1). Examination of the conformational energy landscape was performed for a FSHR model and the structure of the LHCGR (PDB:7FII). Both GPCRs were relaxed in similar conditions through a simulation box that included water molecules as solvent, a polyunsaturated phospholipid membrane, and monovalent ions for charge balance. A difference to highlight between both structures is the presence of the disulfide bond C292-C338 at the HR of the FSHR, which is not observed in the equivalent position (C304-C336) in the LHCGR [19, 20].

In the study of protein folding, it has been proposed the notion of the “energy funnel” that stretches down as the energy decreases [22]. When a protein is unfolded, without a well-defined structure, its possible conformations show high variability that is represented by the top width of the funnel; as the molecular interactions favor the formation of contacts among residues, (*e.g*. hydrogen bonds formation), the free energy of the protein will decrease and the overall, three dimensional shape of the protein (*i.e.* its stable set of conformations) will be better defined, moving towards the lower portions of the funnel. When the protein reaches its native state, the overall conformation will move towards the end of the funnel exhibiting a minimal free energy. The progression of changes from the unfolded to the native (folded) state is rather a rough surface, with local minima and metastable states that should evolve to the more stable conformations. In a given GPCR, as the FSHR, the surface of the conformational energy landscape may show more than one stable state that represent either the inactive state or distinct active, signaling states that may lead to full or biased signaling [23]. In this scenario, knowing the transition route to favor a particular signaling over other(s), represent, on one side, the understanding of the conformational changes occurring during the signal propagation process and, on the other, the possibility to control the selective signaling of a GPCR through different agonists, antagonists or allosteric modulators. The present study explores the energy landscape of the LRR-HR-TM in gonadotropin receptors, to identify the configurations compatible with the opening of their intracellular domains during activation.

From the computational point of view, the identification of critical sites for the allosteric regulation of protein function has been analyzed employing different approaches including machine learning, bioinformartics to detect allosteric sequences, site-directed mutagenesis, and molecular dynamics [24–26]. A widely employed technique to describe conformational changes of a protein is the analysis of the principal component (PC), which consists in the calculation of the covariance matrix whose diagonalization gives rise to a coordinate transformation [27]. The eigenvalues contain the mean square fluctuations associated to each eigenvector, ordered in decreasing order. Through the trajectory projection on eigenvectors, the set of PC discloses the low frequency motions captured in the simulation. Typically, the first PCs accumulate the larger variability with respect to a reference structure (*e.g.* average structure). There are different indicators to establish whether the PC corresponds to sampling with a representative variability of conformational changes, for example, the calculation of the autocorrelation function, the contribution of the thermic fluctuation (content of the temporal periodicity of the cosine function in a PC), and the calculation of the total fluctuation as the square root of the sum of eigenvalues divided by the number of atoms, among others [28]. The calculation of the probability distribution of the first components (e.g. CP1-4) is of particular interest since a projection of the free energy (ΔG) can be calculated form the corresponding histograms in 1 to 3 dimensions. As a result, it is possible to obtain conformations that cluster together as function of the PC values, allowing identification of those conformations with the higher probability as well as the energy barriers associated with a transition. In addition, it is possible to detect intermediate states through which the transition is carried out, which is quite informative for the description of the transition states and the landscape of the conformational energy. Nevertheless, there are effects that may interfere in the analysis: *i.* The adjusting effect to a reference structure (initial, final or average structure) to eliminate translational or rotational displacements; *ii.* The contribution of highly flexible zones that may affect the adjustment with the reference structure; and *iii.* The time scale of the simulation that may not be sufficient for sampling convergence [29, 30]. Some strategies to mitigate these limitations are: *i.* The use of internal coordinates instead of cartesian coordinates, that is, the analysis can be performed using dihedral angles and distance between atoms or domains; *ii.* To restrict the analysis on rigid regions; and *iii.* To carry out bias simulations through the definition of a reaction coordinate (collective variable), implementation of an accelerated sampling method as the replica exchange, trajectories at high temperature, metadynamics, among some [31]. In the present study the exploration of the conformational surface of the GPCRs was performed through a combination of strategies, such as the use of the distances or angles between the LRR, HR and TM, and PC analysis. This study has the perspective of evaluating different coordinates of reaction that allow to capture the transition routes through the intermediates compatible with the conformational switches known to be involved in GPCR activation, namely the D/ERY ionic motif, the displacement of TM helix 6, and the rearrangement of helices 5 and 7 [2, 32]. In addition, a trajectory of the LHCGR in 1-palmitoyl-2-oleoyl-sn-glycero-3-PC (POPC) was generated as the continuation of the initial processing in CHARMM-GUI for further comparisons.

## RESULTS

### Detection of conformational changes from domain distances

Describing the conformational states for the glycoprotein hormone receptors, obtained either by cryo-electron microscopy or computational models, is relevant for understanding of the mechanisms associated with receptor function. In cases when the resolved structure is incomplete, modeling strategies emerge as an alternative approach to unveil processes at the atomic level. To convey the molecular complexity by which membrane receptors link an extracellular stimulus with an intracellular response, in Fig. 1A we show the relaxed structures of the FSHR (AF-P23945) and LHCGR (PDB:7II) in a SDPC membrane environment, identifying the LRR, HR, and TM domains; in addition, Fig. 1B shows the TM domain of FSHR-TM model in comparison to the full FSHR model. From a structural alignment of the helices, the comparison suggests consistent predictions on the relative positions of the TM helices as well as the location of residues, shown by the cysteine residues as an example (Fig 1B). In previous studies, the FSHR-TM model, including D355 to N678, was developed before the breakthrough of the AF2 method for predicting native structures. The good agreement of the FSHR structures obtained provides an additional criterium to validate our former FSHR-TM model [15]. By structural alignment of the LRR of the FSHR (A48-T270) and the LHCGR (T52-T274), a RMSD of 2.40 Å was calculated, whereas for the HR domain of the FSHR (P276-T345) and the LHCGR (P276-P345), the calculated RMSD was 17.9 Å. By aligning the sequence of the TM of the FSHR (Y362-F630) and the LHCGR (D360-T630), we calculated a RMSD of 6.1 Å. The relative displacement between the FSHR and the LHCGR at the intracellular side of helix 6 was 9 Å, which is close to the 14 Å displacement between the reported active and inactive states of the LHCGR [19]. In rhodopsin, the opening of the intracellular sides of TM6, along with rearrangement of helices 5 and 7, favors the open conformation for coupling of G proteins, G protein-coupled receptor kinases (GRK), and arrestins [32]. Therefore, from the position of the TM5 and TM6, which is consistent with an open intercellular conformation, the FSHR model obtained corresponds to the active state. The largest difference between the FSHR and LHCGR was observed in the HR domain, which deserves a detailed analysis because its role on triggering the activation process of these receptors [20, 33]. In this study, we explored the motion of the LRR, HR, and TM consistent with the active state. For this purpose, we defined positions at each domain and calculated the relative distances, which may serve as reaction coordinates to follow the transition among conformational states. In table 1 the atoms selected to measure relative distances of the receptor domains, and their values at the starting conformation are shown. For the LRR we chose a residue at the middle of the first β-strand, for the HR a residue at the middle of the α-helix, and for the TMD a residue at the middle of TM helix 3.

**Table 1.** Relative distances among the LRR, HR, and TM domains of the FSHR and LHCGR.

| Distance | RHLCG | RHEF |
| --- | --- | --- |
| **RLL-BR** | (S55-C304) | (L31-Y303) |
|  | 66 Å | 63 Å |
| **TM-BR** | (L452-C304) | (L460-Y303) |
|  | 32 Å | 56Å |
| **RLL-TM** | (S55-I431) | (L31-L460) |
|  | 32 Å | 56Å |

The comparison among relative distances LRR-TM, LRR-HR, and TM-HR are shown in Fig. 2, including the correlation coefficient for each replicate. The overlapping areas represent conformations with the same values of relative distances (internal coordinates) and larger colored areas represent broader distributions. In R1, distributions were mainly unimodal centered at 70 Å, 85 Å, and 93 Å, for LRR-HR, LRR-TM, and TM-HR distances, respectively. Positive correlations calculated for distances LRR-TM and TM-HR were of 0.40, 0.55, and 0.4, for R1, R2, and R3, respectively (Fig. 2C). The increase of the LRR and HR distance relative to TM was consistent with a shortening of the LRR and HR distance, as can be observed in R1 by the negative correlation between LRR-HR and LRR-TM. From the distributions of relative distances among the LRR, HR, and TM domains, covering broader intervals (*e.g*. 80 to 100 Å between TM and HR, or 72 to 93 Å between LRR and TM (Fig 2C), they can be used as reaction coordinates to identify conformational intermediates. Interestingly, in replicate R3 the distance variability was smaller than in replicates R1 and R2, which can be related to the sampling of a metastable state. Metastable states may prevent transitions among intermediaries and alternative strategies must be implemented for an extensive sampling of the conformational space. For example, trajectories can be generated with configurations harvested from a previous run, as it is described below.

**Fig. 2.**
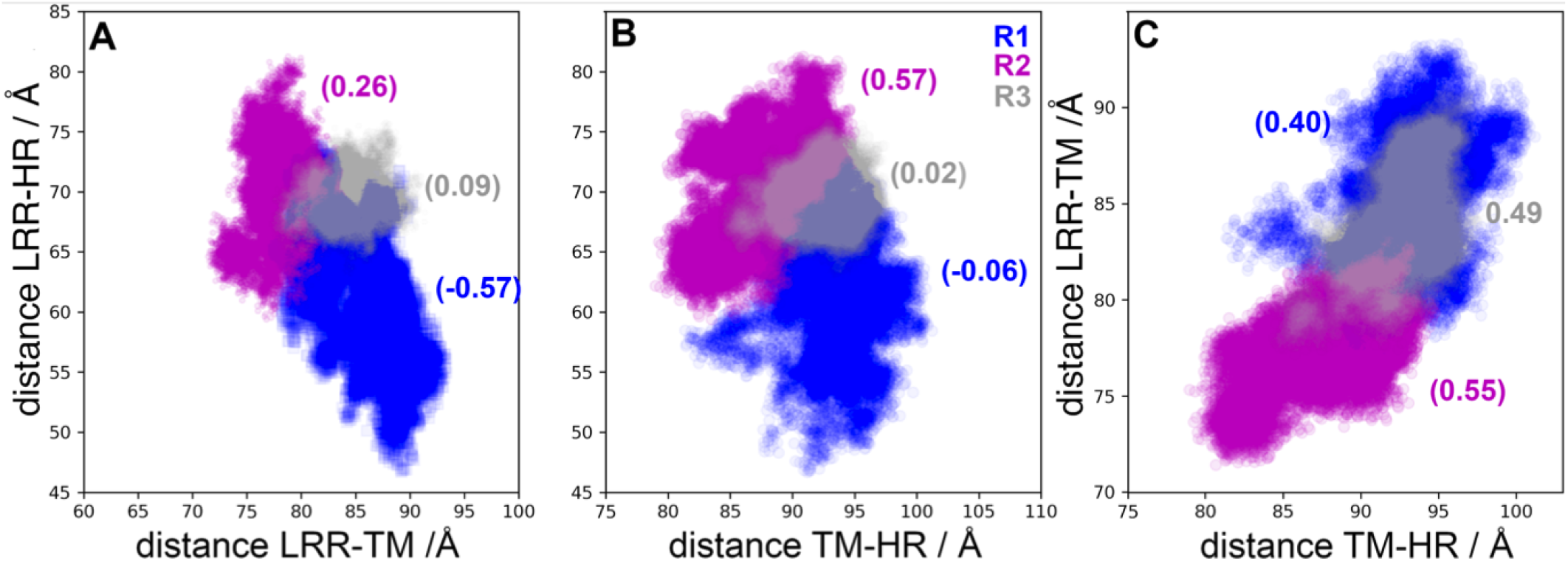
Distances among structural domains of the FSHR. Numbers in parenthesis indicate the correlation coefficients for every pair of distances. Color code: R1-blue, R2- magenta y R3-gray.

The correlations for distances in LHCGR are shown in Fig. 3 for replicates R1-3 and for the trajectory in POPC. From the trajectory in POPC, receptor configurations were harvested at t=0 ns (R1), t=100 ns (R2), and t =180 ns (R3). Given the initial configurations of R2 and R3, these trajectories displayed no overlaps with replicate R1. Instead, the trajectory in POPC connected the explored regions by R2 and R3 with R1. Interestingly, the trajectory in POPC showed multimodal and broader distributions than replicates R1-3 for distances LRR-HR and TM-HR (Fig. 3B). All trajectories consistently showed negative correlations between LRR-HR *vs* TM-HR (Fig. 3B), which indicated that the HR was moving away from the LRR, approaching to TM. Correlations of distances LRR-TM *vs* TM-HR in the LHCGR (Fig. 3C) showed different trends than those in the FSHR (Fig. 2C); the concerted motion of the LRR and HR relative to TMD in FSHR was not detected in the LHCGR. In summary, from the sampling of receptor conformations in independent replicate trajectories, starting from the same initial configuration (with restarted velocities) or from conformations harvested from a previous trajectory, it was possible to prevent sampling of metastable states. In fact, the weighted ensemble (WE) method takes advantage of generating multiple, short trajectories departing from different initial configurations and restarting trajectories, whenever the reaction coordinate populates a new bin along the state’s transition [34]. In Fig. S1 of SI_info, it is shown a plot of the RMSD for the active state of the LHCGR obtained from 100 iterations implementation of the WE method. The RMSD can be used, in fact, as reaction coordinate to track the transition between the active to inactive states.

**Fig. 3.**
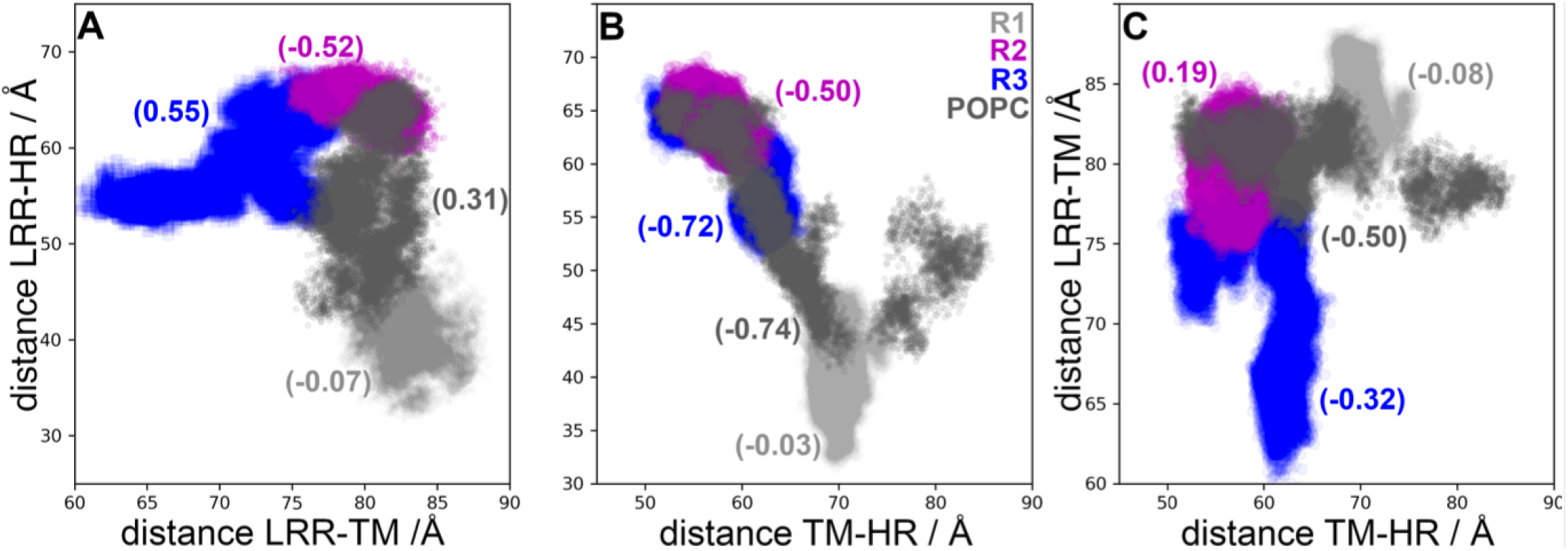
Distances between structural domains of LHCGR. Numbers in parenthesis are the correlation coefficients for every pair of distances. Color code: R1 gray, R2 magenta, and R3 blue, POPC carbon gray.

### Conformational analysis of LHCGR

Fig. 4 shows RMSD matrices for the LHCGR replicates R1-3 and the POPC; insets also show cluster conformational analysis with a cut-off criterium RMSD <0.5 Å. For the RMSD calculation, only the TM helices were fitted in time frames at *t* and *t+Δt*: TM1, 360-387; TM2 392-420; TM3, T437-470; TM4, 480-504; TM5, 522-552; TM6, 565-597; and TM7, 602-625. The red solid line in the plots was drawn to identify self-similar groups. Not surprising, similar structures were found along the diagonal as conformations in consecutive frames differed within the cutoff criterium. RMSD values in the TM1-7 helices were lower than 3.1 Å in SDPC and lower than 2.8 Å in POPC. In Fig. 4A, self-similar groups were detected at the ∼10 ns time scale in R1. In R2, a group was detected in the first 50 ns and other after 100 ns (Fig. 4B). In R3, RMSD values showed closer differences in comparison with R1 and R2. In the cluster analysis, a structure was added whenever it showed a RMSD within the cut-off of any of the members in that cluster. For example, with a cut-off of 1 Å only one cluster was detected; therefore, we set a 0.5 Å for detection of larger number of clusters. Clusters were calculated for each replicate in SDPC and the POPC trajectory, altogether encompassing ∼800 ns of conformational sampling.

**Fig. 4.**
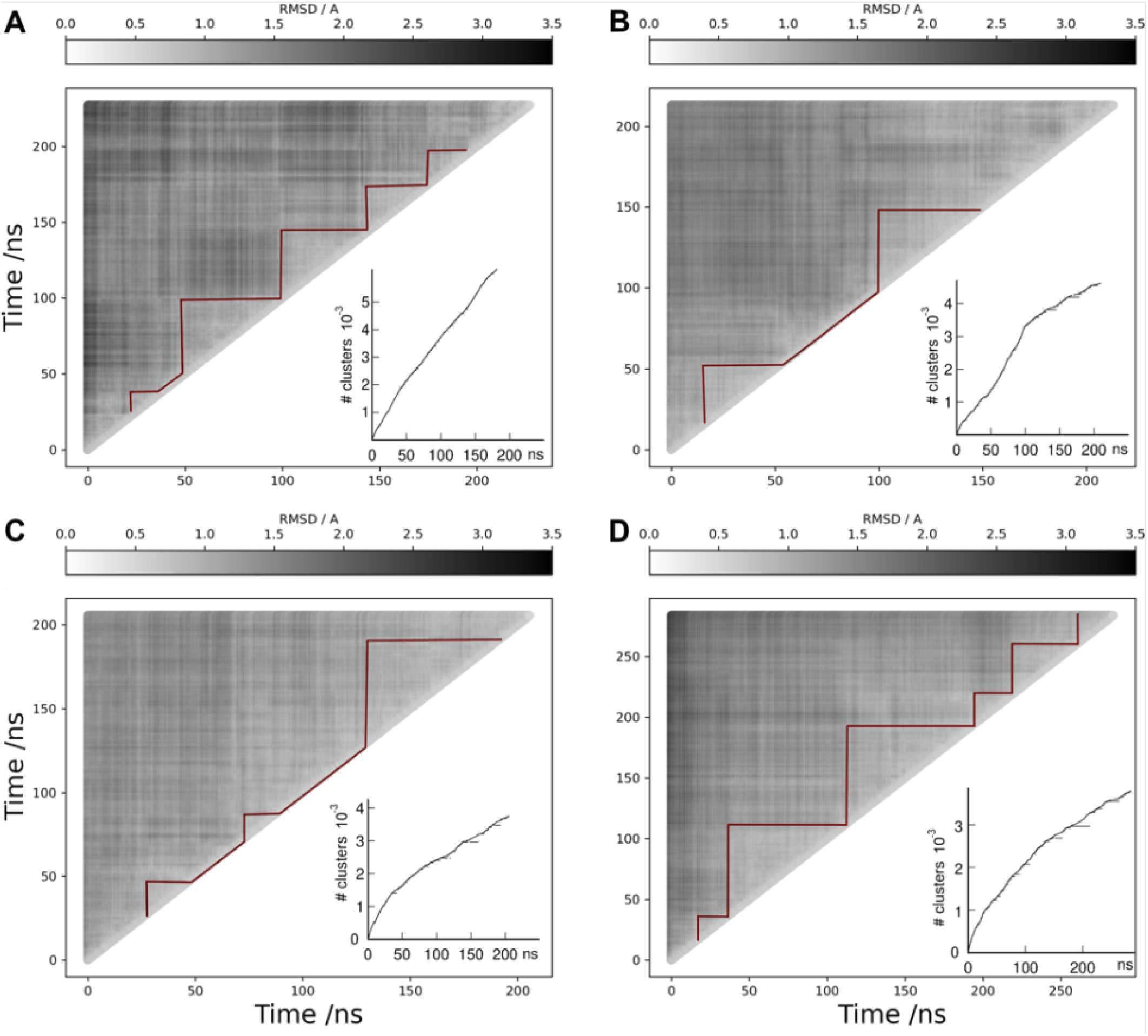
RMSD matrix analysis (gray scale) for the TM helices of LHCGR. (A) replicate R1, (B) replicate R2, (C) replicate R3, and (D) trajectory in POPC. Red solid lines along the diagonal were drawn for visual identification of self-similar groups. The cluster analysis is shown in the insets. Conformations in clusters include structures within 0.5 Å of RMSD among each other. The cluster number is identified in the vertical axis. Horizontal segments along the curve identify the time frames forming a given cluster.

PC analysis for the LHCGR runs is shown in Fig. 5 by means of 2D projections of replicates R1-3 over the first four eigenvectors. For the largest clusters found in each replicate, the set of conformations were also projected over the eigenvectors: cluster 6383 of R1, clusters 4193 and 2595 of R2, and clusters 1404 and 2964 of R3. Conformations in clusters were plotted in the context of the full trajectory (Fig 5). Further, in Fig. 6 the distributions for each PC of the replicates are shown, providing an additional context in which the clusters were located. For example, conformations in cluster 6383 of R1 were not at the maximum of -8 nm, but instead they were located at 0 nm near of the tail of the distribution (Fig 5A). In R2, the PC1 varied from -50 nm to 10 nm, with maxima at -30 nm, -8 nm, and 8 nm; cluster 4193 contributed with conformations at the 8 nm maxima, whereas cluster 2595 did so with conformations at -30 nm. Cluster 2964 of R3 had conformations at the maximum of 2.3 nm of PC1; cluster 1404, around the maximum of 22 nm (Figs. 5C, 6A). Because of the broad shape of the PC3 of R1 (Fig 6C), cluster 6383 showed values that spread from -5 nm to 5 nm, but right on maximum of PC4 at -2.7 nm (Fig. 6D). Cluster 4193 of R2 showed a population at the maxima of PC3 (-5.4 nm) and PC4 (-4.0 nm); and cluster 2595 at 7.5 nm of PC4 (Figs. 5E, 6D). Finally, clusters 2964 and 1404 of R3 (Figs. 5F, 6C) had conformations at maxima of PC3 (3.8 nm) and PC4 (3.3 nm; Fig 6D). By combining cluster and PC analysis for the TM conformations, we could identify those conformations that most likely contributed to the fluctuations of the low frequency motions, that is, the *intermediary states* in a free energy landscape.

**Fig. 5.**
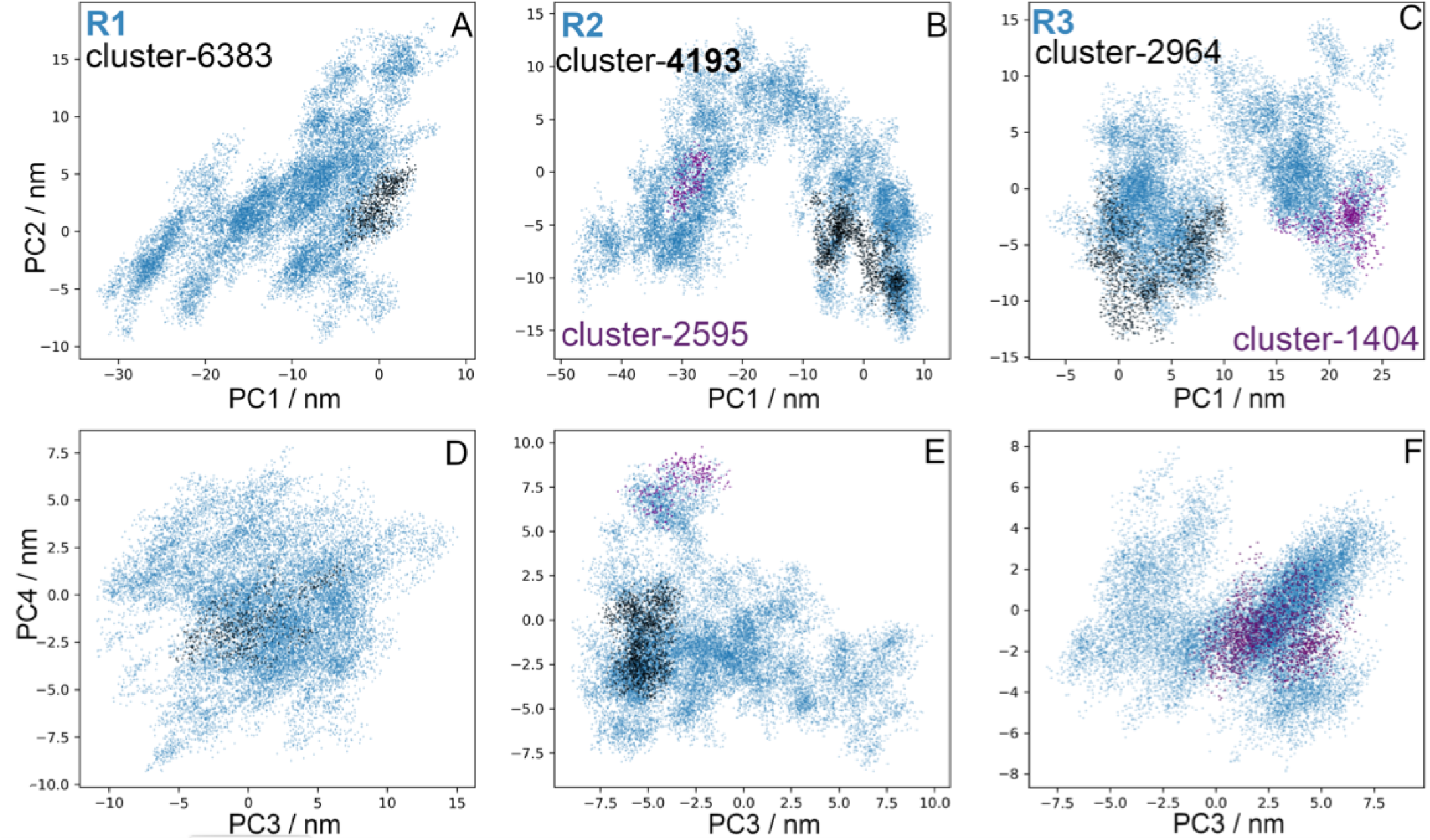
Principal component analysis for LHCGR. PC1 and PC2 (A-C) and PC3-PC4 (D-F) of trajectories R1-3 (blue dots). Projections for cluster 6383 of R1 (A, D), 4193 of R2 (B,E) and 2964 of R3 (C,F; black dots); clusters 2595 of R2 (B,E), and 1404 of R3 (C,F; purple points). TM domain conformations in clusters were identified with a RMSD <0.5 Å, *ie.* a structure is included whenever its RMSD is lower than 0.5 Å from any of the members in that cluster.

**Fig. 6.**
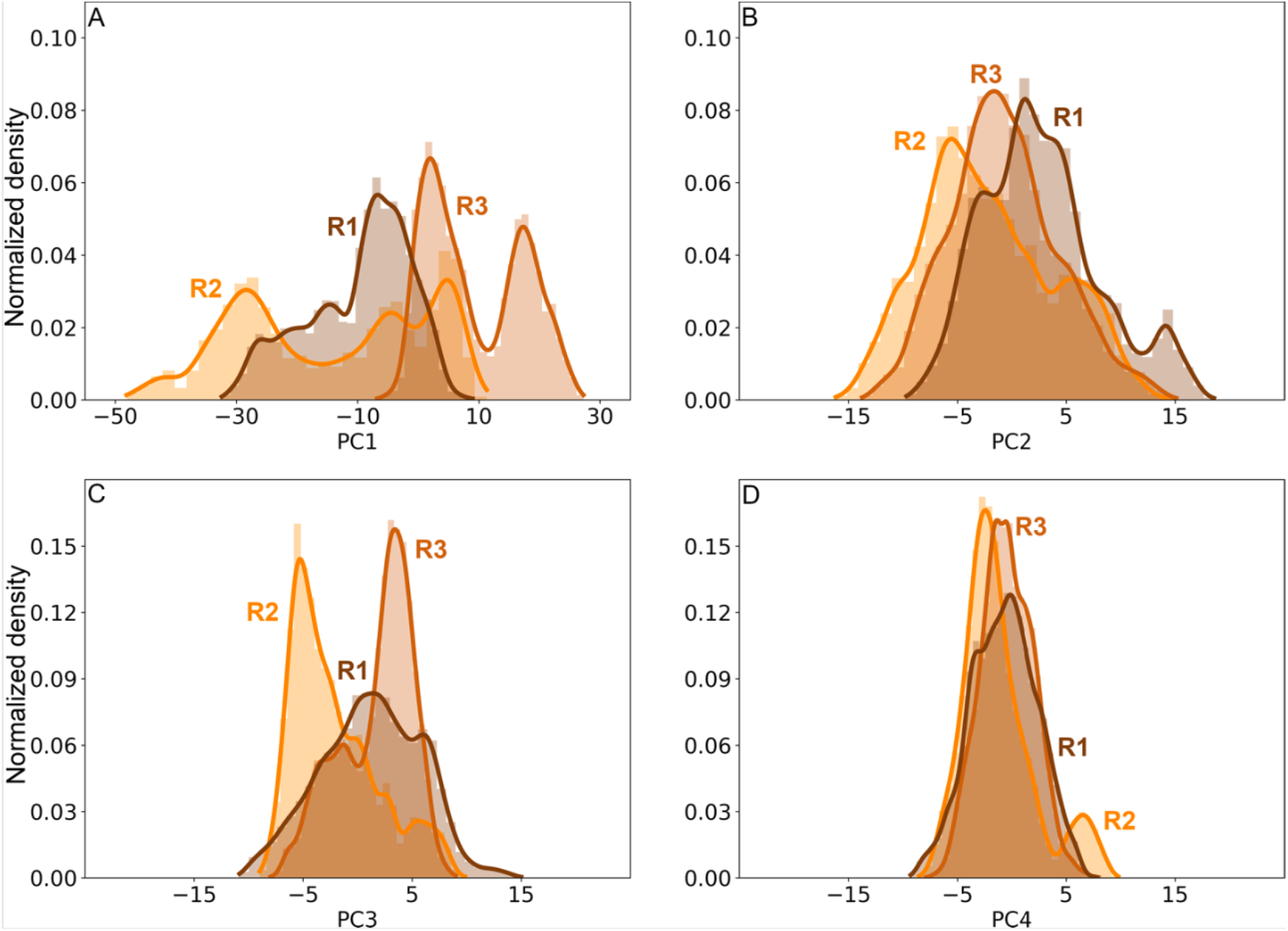
Distributions of the trajectory projections on the first four eigenvectors for the LHCGR in replicates R1 (brown), R2 (orange), and R3 (brown-orange). PC1 (A), PC2 (B), PC3 (C), and PC4 (D).

### Conformational analysis of FSHR

The motion of the receptor LRR and HR domains detected by the relative distances, can be related to the fluctuations of the TM as a correspondence between the dynamics of the aqueous and membrane environments. It seems evident that by including the variability of the LRR and HR domains together with the fluctuations of the TM helices in a conformational analysis, makes difficult to identify the transition states due to the slightness of helices motions [32]. In Fig 7 the RMSD matrices and the cluster analysis for replicates R1-3 of the FSHR are shown. Only the TM helices were fitted for structures taken at time t and t +Δt: TM1, Y362 to Y392; TM2, P397 to Y432; TM3, N437 to T472; TM4, A487 to G507; TM5, M532 to R557; TM6, D567 to L597; and TM7, A607 to Y626. Calculated values of RMSD were lower than 2.5 Å, 2.2 Å and 1.9 Å, for R1, R2, and R3, respectively. The cluster analysis is included in insets of Fig. 7, using a RMSD cut-off <0.5 Å; a structure is included whenever the RMSD was <0.5 Å in any of the members of that cluster.

**Fig. 7.**
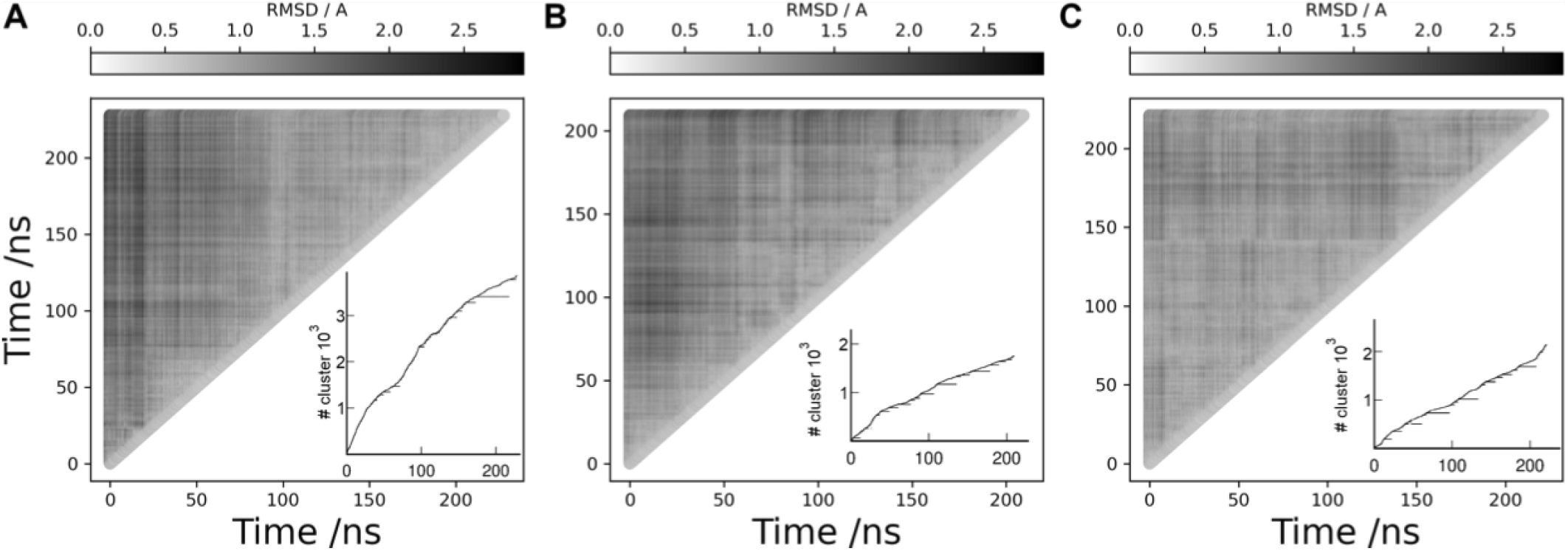
RMSD matrix analysis (gray scale) for the TM helices of FSHR. **A**. replicate R1, **B**. replicate R2, and **C**. replicate R3. The cluster analysis is shown in the insets. Conformations in clusters include structures within 0.5 Å of RMSD among each other. The cluster number is identified in the vertical axis. Horizontal segments along the curve identify the time frames forming a given cluster.

The 2D projections of replicates R1-3 on the first four eigenvectors are shown in Fig. 8: PC1 *vs* PC2 and PC3 *vs* PC4. The RMSF of PC1 represents ∼70% of total fluctuation, whereas the RMSF of PC1 to PC4 ∼95% (Fig. S2 in SI_info). Clusters 3405, 1171, and 0733 were identified for R1, R2, and R3, respectively, with the largest number of conformations within the RMSD cut-off. To identify the most populated intermediaries in the context of the full trajectory, projections of the clusters on eigenvectors were included in Fig. 8. In addition, Fig. 9 shows the distribution of the PC1 to PC4 for each replicate. Cluster 3405 of R1 showed values of -0.5 Å in both PC1 and PC2 (Fig 8A), which was consistent with the maxima of their distributions (Figs. 9A, 9B). Due to the broad distributions of PC3 and PC4 (Figs 9C and D), conformations in cluster 3405 were as disperse as the full trajectory. Cluster 1171 of R2 showed conformations around the maxima of PC1 to PC4 (Figs. 8E and F), according to its corresponding distributions (Figs. 9C and D). Identification of conformations at the maxima of the main PC may underpin an important intermediary. In R3, distribution of PC1 displayed a prominent maximum at -16 nm and a secondary maximum at -25 nm (Fig 9A); PC2 showed a broad distribution with four maxima from -20 nm to 15 nm (Fig. 9B). Cluster 0733 of R3 showed a conformation at maxima of all PC, albeit not in the principal maximum of PC3. In summary, the most populated clusters whose conformations of the TM domain exhibited a RMSD <0.5 Å, populated the PC distributions with conformations showing high probabilities. By identifying less populated clusters, the intermediates in values at shoulders or minimum of the PC distributions could be useful to establish possible transition states.

**Fig. 8.**
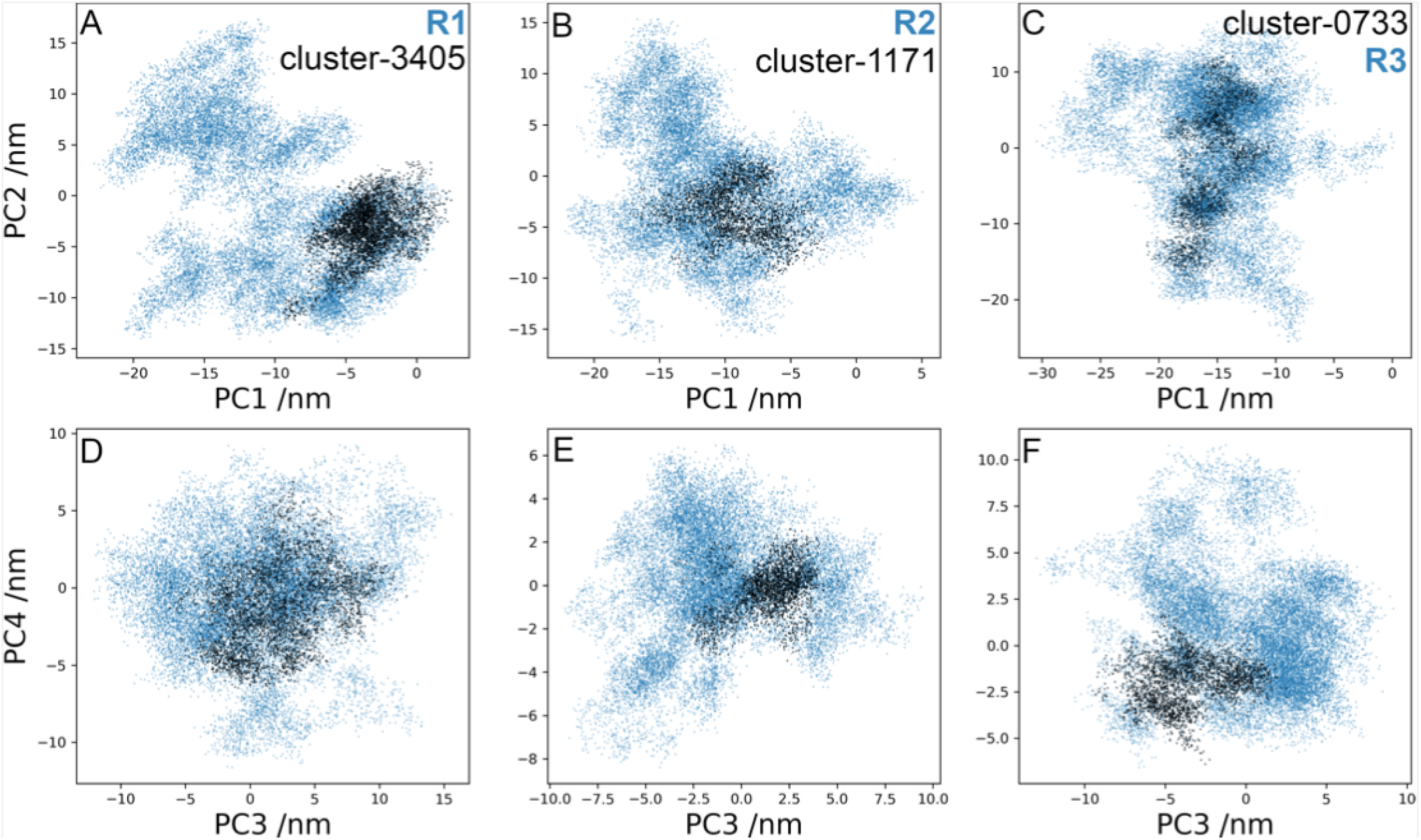
Principal component (PC) analysis in the FSHR. Projections on PC1-PC2 (A-C) y PC3-PC4 (D-F) of trajectories R1-3 (blue points). Projections for groups 3405 in R12, 1171 in R2, and 0733 in R3 are included (black points). The groups conform subclusters of conformational states whose RMSD difference in the TMD region is <0.5 Å.

**Fig. 9.**
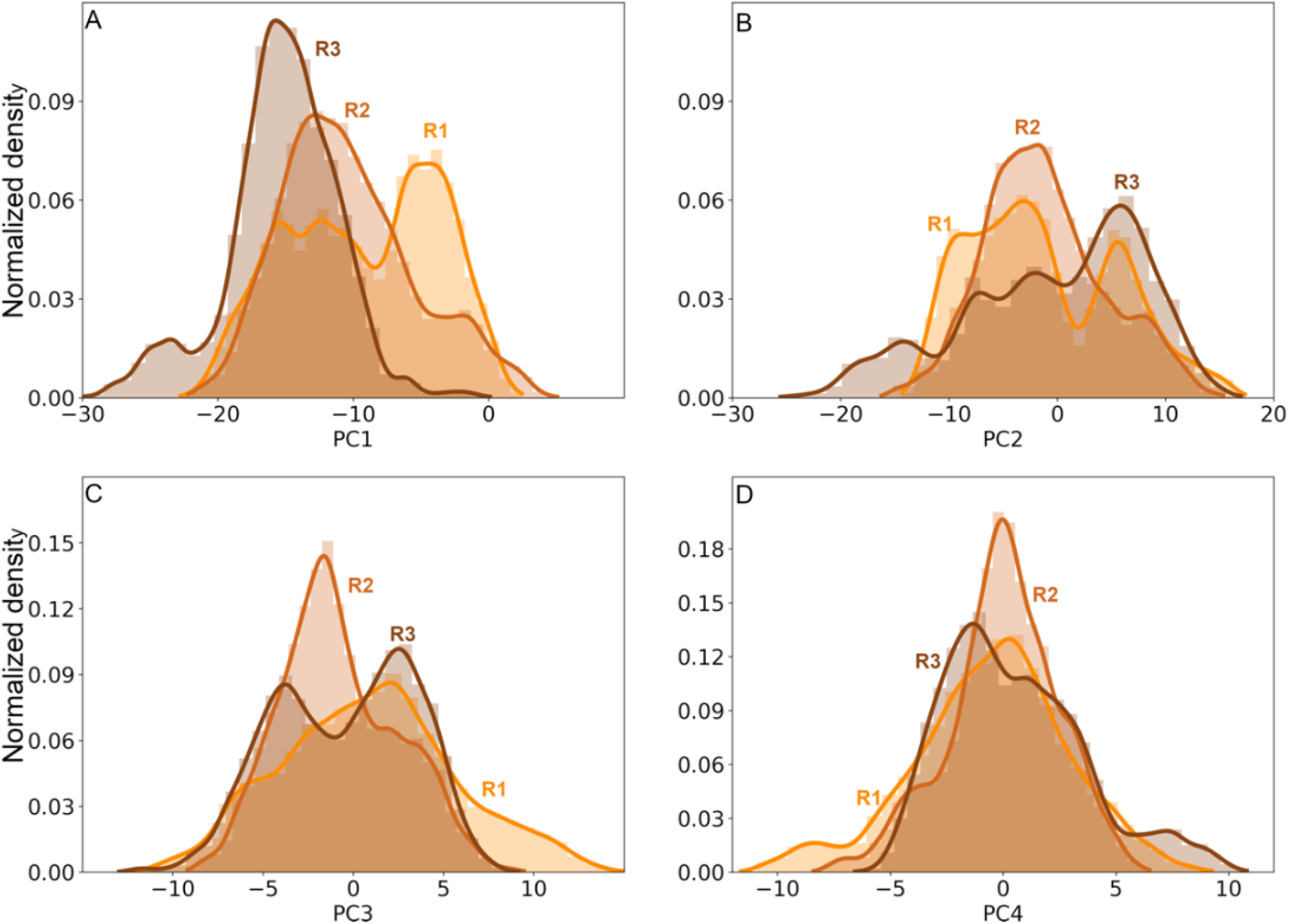
Distributions of the trajectory projections on the first four eigenvectors for the FSHR in replicates R1 (orange), R2 (brown-orange), and R3 (brown). PC1 (A), PC2 (B), PC3 (C), and PC4 (D).

### Correspondence between membrane and aqueous dynamics of the receptor’s domains

Clusters conformed by the cut-off criterium (RMSD <0.5 Å) may be related to motions of the extracellular domains, LRR, HR, and TM. Fig. 10 shows distances for the clusters identified in replicates R1-3 of the LHCGR. Cluster 6383 was detected for R1 (Fig. 10, row R1); however, the cut-off criterium produced too many clusters with few conformations (Fig. 4A). To circumvent this problem, the cut-off can be adjusted to values 0.5 <RMSD< 1.0 in order to conform more populated clusters. In fact, for R2 the RMSD cut-off criterium allowed to identify distinct clusters in terms of the relative LRR, HR and TM distances (Fig. 10, row R2); hence, each cluster corresponds to a different state of relative distances. In R3, there were also distinguishable clusters, albeit in this case there were overlapping conformations most likely because clusters 2407 and 2964 were close in time (Fig. 10, row R3). In addition, in R3 it was possible to detect the negative correlation between RRL-RB *vs* TM-RB showed in Fig. 3B. From the analysis of relative distances in clusters, it was possible to identify conformational states of the LHCGR in MD trajectories for independent replicates, with starting configurations at different times of a previous run. Using both, the RMSD and any of the relative distances, it is possible to follow transitions among conformational states (*e.g.*, active—inactive) with these reaction coordinates.

**Fig. 10.**
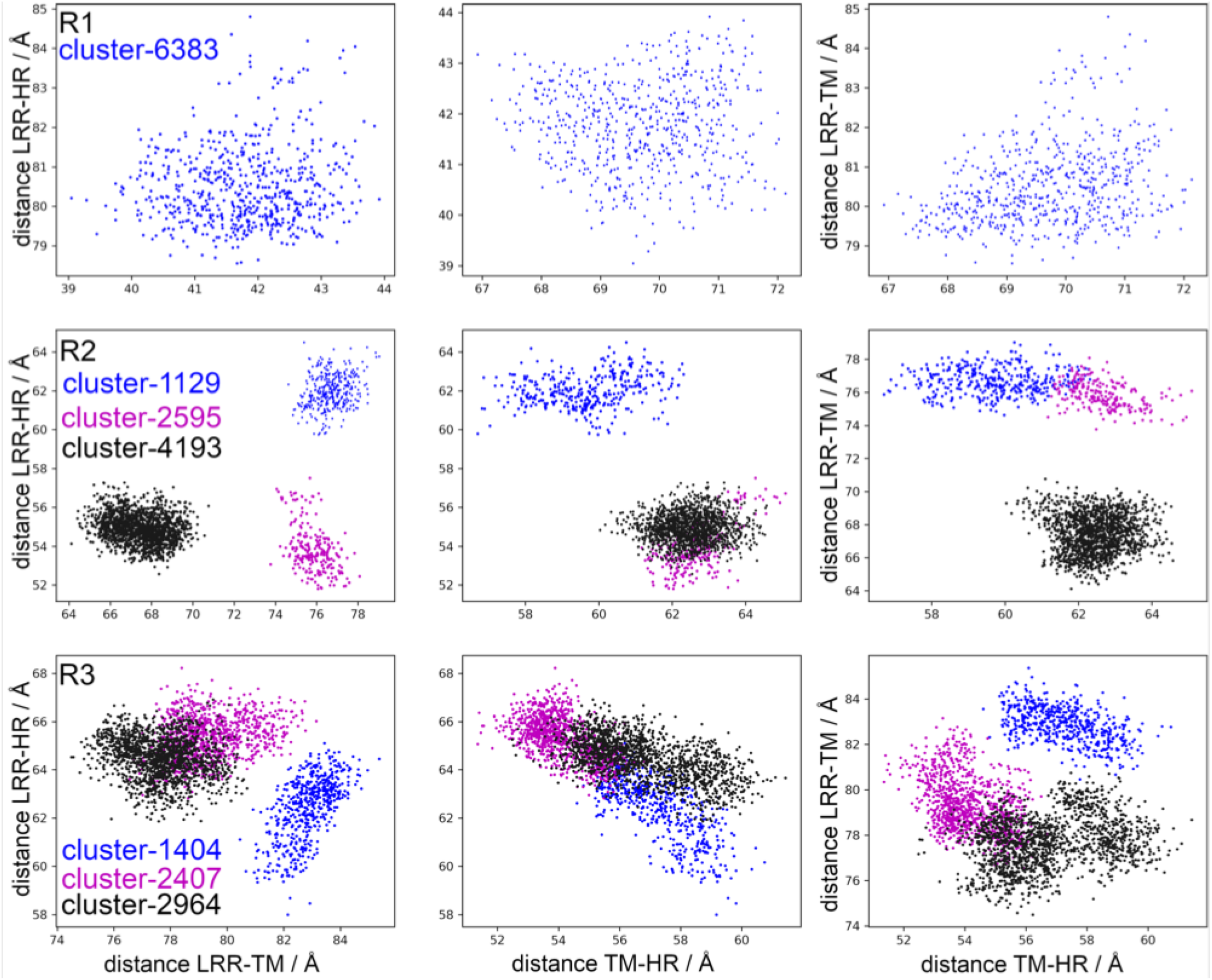
Distances among the LRR, HR and TMD in the LHCGR (Å) in R1 (upper panel), R2 (middle panel) and R3 (lower panel). Analysis for clusters 6383 of R1, 1129, 2596, and 4193 of R2, and 1404, 2407, and 2964 of R3.

Figure 11 shows the relative distances for the FSHR and, for this particular case, for only the most populated cluster of each replicate R1-3. Conformational states can be identified in terms of the relative distances: in particular, R2 showed almost no overlapping conformations with R1 and R3. Because the trajectories were independent, the clusters explored different regions of the conformational energy landscape. More comparisons can be made among replicates; for example, it is possible to project a trajectory over the eigenvectors of a second trajectory. In Fig. S3 of SI_info, projection of R1 over the first eigenvector of R2 provides information on the conformations of R1 that contributes to PC1 of R2; conversely, a lack of overlap among distributions would represent that trajectories show different dynamics in the conformational space. By the strategy applied in this study, we could distinguish conformations of the FSHR in which the HR moved relative to LRR, as suggested by the previously proposed activation mechanism [7].

**Fig. 11.**
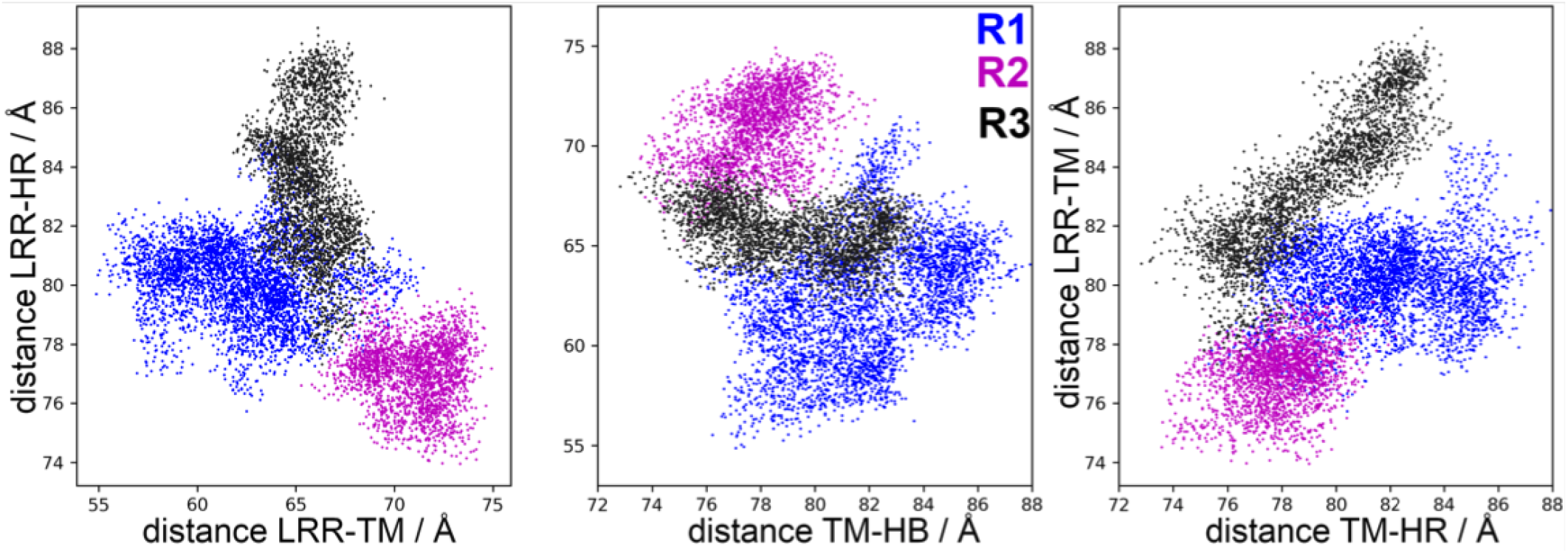
Distances among LRR, HR and TM domains for the FSHR (Å). Blue dots, group 3406 of R1; magenta dots, group 1171 of R2; and black dots, group 0733 of R3.

## DISCUSSION

In this study we explored the conformational changes of the gonadotropin receptors (subfamily of GPHR), in which the FSHR is the prototypical member, within the family A of GPCR. One of the key physiological functions of GPCRs is signal transduction, that is, the triggering of a cell response to a particular (agonist) stimulus. Nevertheless, these receptors are not necessarily a switch that turns on and off in response to a stimulus, but rather exhibit diverse responses [18]. Conformational variability might explain that a particular receptor may be coupled to distinct signaling molecules, which in turn is related with different concepts such as allosteric regulation, signal predisposition or selective signaling [35, 36]. Given that the structural information to determine the intermediate states during the activation process of GPCRs still is scarce; in silico molecular dynamics techniques are quite useful to establish their structure-function relationship in such dissimilar environs, from the phospholipid membrane core to the bulk aqueous medium. In particular, we employed a FSHR model that included the LRR, HR and TM as it is reported in the AF2 server [37, 38]; the LHCGR structure was obtained from the PDB repository (access code 7FII) [19]. GPHRs were analyzed in a comparative manner following the same computational protocols. In both systems, the internal coordinates were defined to measure distances among the LRR, HR and TM domains to detect the relative motion of the HR, which is recognized as a key region involved in the activation of these receptors [20, 33, 39].

The HR exhibits an α-helix segment, a P10 segment and a loop which extends to the aqueous medium that resembles the thumb of a glove (Fig. 1). The activation mechanisms proposed based on the LHCGR structure in the active state, consists in the displacement of the LRR in vertical position with respect to the plasma membrane (the TM-LRR distance increases) while the HR approximates to the membrane (TM-HR distance decreases) [19]. In the LHCGR, in the present generated trajectories exhibited relative motion in which the LRR-TM fluctuates between 59.7 and 88.6 Å, the LRR-HR between 31.4 and 70.0 Å, and the TM-HR between 48.2 and 79.2 Å. Motions of the TM- HR and LRR-HR consistently exhibit negative correlations, where the increase in the former tends to decrease the latter. In the crystal structure, the presence of the hormone of the LHCGR may prevent the proximity between the LRR and the HR. These extreme values may be useful to implement weighted ensemble simulations [34].

Although the role of the HR as an activation switch in GPCRs like the FSHR has been recently challenged [39], it has been proposed that during FSHR activation a displacement of the HR that allows insertion of residue Y355 in the interface between the α and β subunits of FSH occurs [7, 40]. In the active state, the LRR is detected in vertical position (perpendicular to the membrane) and the HR must move to the LRR [19, 41]. That is, the HR movement consists in an increase in LRR-TM distance and a corresponding decrease in the LRR-HR one, leading to a negative correlation between those distances. In the FSHR the relative motions of the LRR-TM would fluctuate between 71.4 and 93.3 Å, those of the LRR-HR between 46.7 and 81.3 Å, and those of the TM- HR between 78.9 and 101.2 Å. Hence, trajectories using configurations with increasing TM-LRR and/or decreasing LRR-HR distances, along with fluctuations of RMSD at the TM domain, could be useful to identify transition intermediaries.

Another important element associated to the motion of the HR is the rearrangement of the EC2, which has an inhibitory effect on the TM. In our simulations, this effect on the TM helices was not observed because its initial configuration was already in the active position, with the intracellular portions of the helices 5 and 6 displaced out. Nevertheless, by means of a *distance* criterion (RMSD <0.5 Å), clusters showed well differentiated conformations in the extracellular domains, in particular, the relative position of the HR. In the present study, the extracellular and TM domains were initially studied uncoupled due to the difference in the dynamics of the aqueous media and the membrane; thereafter, it was possible to identify TM clusters with specific values of relative positions of the LRR and the HR. Therefore, to explore the transitional states between the active — inactive states, we propose that LRR-TM, or TM-HR relative distances may be useful as reaction coordinates.

The purpose of exploring the energy landscape of the gonadotropin receptors is, among others, to screen for structural variations that may respond to conformational changes leading to well-known states, or alternative states; for example, those promoting binding of new drugs, or selective coupling to intracellular transducers. A broad perspective could include using the reactions coordinates, here explored, to bias the conformational dynamics of the receptors upon binding allosteric modulators or agonists able to favor particular signaling pathways, as required by a given therapeutic purpose.

## MATERIAL AND METHODS

Two simulation boxes containing the receptor, a lipid bilayer, water molecules as solvent, and Na^+^ or Cl^-^ ions for charge balance were set up. For the FSHR initial coordinates, we used the AF2 [37, 38] structure, generated with the full sequence of the human receptor model (AF-P23945). Post-translational modifications were introduced at C644 and C646 by palmitoylation through thioester bonds [21]. Disulfide bonds between the cysteine side chains of C18-C25, C23-C32, C275-C346, C276-C356, C442-C517, and C229-C338 were defined according to the S-S bond distance criterium. Protonation states of side chains were set to those of the predominant species at neutral pH. The FSHR structure was inserted in a preequilbrated bilayer of 1-stearoyl-2-docosahexaenoyl-sn-glycero-3- phosphocholine (SDPC). Water molecules for solvation of the receptor and the lipid heads were added in a rectangular box of 120X120X160 Å^3^, respectively, in the x, y and z directions. A total of 191601 atoms were included: 48411 water molecules, 281 lipids, 3 Na^+^ ions and the FSHR with 695 residues. The second simulation box contained 190072 atoms: 48414 water molecules, 247 SDPC lipids, 3 Na^+^ ions and 7 Cl^-^ ions, and the LHCGR with 613 residues. The LHCGR structure (PDB:7FII) corresponds to the state bound to the hormone and the G_s_-protein [19], and it was processed according to the CHARMM-GUI membrane builder [42, 43] using the following parameters [44]. The chain R (segment PROD) of the pdb file was selected to be inserted in a POPC lipid bilayer, with box dimensions of 100x100x168 Å^3^. For the missing residues T287 to W329 of the HR, the CHARMM-GUI modeling scheme included the coordinates as predicted by the GalaxyFill algorithm [45]. Disulfide bonds were defined between cysteines C279-C343, C280-C353, C131-C156 and C439-C514. The LHCGR principal axis was aligned in the *z* direction and displaced until the TM domains matched the hydrophobic core of the bilayer. The system size was 191601 atoms: 248 1-palmitoyl-2-oleoyl-sn-glycero-3- phosphocholine (POPC) lipids, 37479 water molecules, Na^+^Cl^-^ ions (0.15 M), and the LHCGR. From the assembly of the initial configuration, the systems were energy minimized with conjugated gradient algorithm for 10 k steps, followed by a gradual relaxation of the receptor atoms. In a first stage, the receptor backbone atoms were fixed, then subsequent stages with positional constrains of 20, 15, 10, 5, 3, 2, and 1 kcal /mol Å^2^, in short trajectories of 200 ps each were applied. The trajectory of LHCGR in POCP was prolonged for 200 ns without constraints. Because of our interest in exploring the receptor’s conformational landscapes, we generated independent trajectory replicates for both the FSHR and the LHCGR in SDPC. In the case of the FSHR, three replicates were generated using the last configuration after the relaxation procedure and restarted velocities for each replicate. For the LHCGR replicates, we used configurations taken from the trajectory in POCP at times 0 ns, 100 ns, and 180 ns, and relaxed the receptor in the membrane environment of SDPC lipids.

All simulations were performed with the NAMD 2.14 software [46], version NAMD3.0 alpha, which was optimized for GPU-accelerated servers [47]. Simulation trajectories were generated in the isothermal-isobaric ensemble (NPT) with Langevin dynamics to maintain a constant temperature [46], and Nosé-Hoover Langevin piston to maintain a constant pressure of 1 bar [48]. Anisotropic cell fluctuations in the x-, y- and z-axis were allowed [49]. Non-bonding interactions were calculated with a cutoff of 11 Å, and a shifting function starting at 10.0 Å. A multiple time step integration for solving the motion equations was used with one step for bonding interaction and short-range nonbonding interactions, and two steps for electrostatic forces, with 2 fs time step. All hydrogen atoms were fixed using the SHAKE and RATLLE algorithms [50, 51]. Electrostatic interactions were evaluated using PME [52], with a 4^th^ order interpolation on a grid of ∼1 Å in the *x-*, *y-* and *z-*directions, and a tolerance of 10^-6^ for the direct evaluation of the real part of the Ewald sum. CHARMM36 all-atom force field parameters were used for the lipid molecules [53], and CHARMM36m for the protein atoms [54, 55], including CMAP correction [56, 57]. Water molecules were modeled using the TIP3P potential [58].

Trajectory analysis were performed in the TCL environment of VMD [59] for the calculation of the root mean square deviation (RMSD), root mean square fluctuations (RMSF), LRR-HR-TMD distances, among other structural and dynamical parameters. For the principal component analysis, GROMACS 2020 [60] was employed with commands *gmx covar* and *anaeig* for the calculation of covariance matrix and analysis of eigenvectors, respectively,. For the first four eigenvectors we calculated the principal components of the C_α_ atoms at the TM helices of the FSHR: TM1, Y362 to Y392; TM2, P397 to Y432; TM3, N437 to T472; TM4, A487 to G507; TM5, M532 to R557; TM6, D567 to L597; and TM7, A607 to Y626; and the LHCGR: TM1, 360-387; TM2 392-420; TM3, T437-470; TM4, 480-504; TM5, 522-552; TM6, 565-597; and TM7, 602-625. RMSD matrices for comparison of frames at *t* and *t+Δt* were calculated for TM helices. Correlations among domain distances also were calculated to detect concerted motions. Visual molecular dynamics (VMD) was used for visualization, representation of 3D structures and image generation [59].

## ACKNOWLEDGMENTS

This study was supported by grant IN208323 from the Programa de Apoyo a Proyectos de Investigación e Innovación Tecnológica (PAPIIT), UNAM, Mexico (to A.U.-A).

